# Ligand Sensing Enhances Bacterial Flagellar Motor Output via Stator Recruitment

**DOI:** 10.1101/2020.04.20.050906

**Authors:** Farha Naaz, Megha Agrawal, Soumyadeep Chakraborty, Mahesh S. Tirumkudulu, K.V. Venkatesh

**Author notes:** Author contributions: F.N, M.A., S.C., M.S.T. and K.V.V. designed research; F.N., M.A. and S.C. performed research; F.N., M.A., M.S.T. and K.V.V analyzed data; and F.N., M.A., M.S.T. and K.V.V. wrote the paper. Equal contribution (Farha Naaz, Megha Agrawal).

## Abstract

The phenomenon of chemotaxis in bacteria, where the cells migrate towards or away from chemicals, has been extensively studied in the past. For flagellated bacteria such as *Escherichia coli*, a change in chemical concentration in its environment is sensed by a chemoreceptor and communicated via a well-characterised signalling pathway to the flagellar motor. It has been widely accepted that the signals change the rotation bias of the motor without influencing the motor speed. Here, we present results to the contrary and show that the bacteria is also capable of modulating motor speed on merely sensing a ligand. Step changes in concentration of non-metabolisable ligand cause temporary recruitment of stators leading to a momentary increase in motor speeds. For metabolisable ligand, the combined effect of sensing and metabolism leads to higher motor speeds for longer durations. Swimming speeds measured at the population level corroborate the observations. Experiments performed with mutant strains delineate the role of metabolism and sensing in the modulation of motor speed and show how speed changes along with changes in bias can significantly enhance bacteria’s response to changes in its environment.

## Introduction

It is well known that bacteria can execute a net directed migration in response to external chemical cues, a phenomenon termed as chemotaxis [1, 2]. One of the most commonly studied bacteria for understanding chemotaxis is *Escherichia coli* [3]. On sensing a ligand, these bacteria transmit the signal from the environment via a well characterised signalling pathway to reversible flagellar motors that drive extended helical appendages called flagella thereby achieving locomotion [4, 5]. The motor is powered by a proton flux, termed as proton motive force (PMF), through an electro-chemical gradient of protons across the inner cytoplasmic membrane [6, 7]. When all flagella rotate in the counterclockwise (CCW) direction (as seen from the flagellar end), the flagella collect into a single bundle leading to a forward motion called a run, whereas when one or more of the flagella rotate in the clockwise (CW) direction, they separate from the bundle leading to an abrupt change in direction, also known as a tumble [8]. By modulating the duration of runs and tumbles, the bacteria achieve a net motion toward chemoattractants or away from chemorepellents [9].

The direction of rotation of the flagellar motor is controlled by the switch complex comprising of proteins FliG, FliM and FliN, which are present at the cytoplasmic end of the motor (Figure 1) [10]. When CheY-P, the phosphorylated state of the chemotaxis protein CheY, binds to the motor protein FliM, the motor is biased in the CW direction [11]. The null state of the motor, however, is the CCW direction. When a ligand binds to a chemoreceptor protein present in the periplasm, the activity of the chemotaxis pathway protein CheA is reduced, which in turn reduces the production of phosphorylated CheY-P. Since fewer CheY-P molecules are available to bind to the motor protein FliM, the flagella rotate in the CCW direction [12]. For ligand concentrations in the nanomolar to the micromolar range, a feedback control mechanism intrinsic to the signalling network brings back the phosphorylation state of CheA and CheY to the prestimulus levels and resets the duration of CW and CCW rotations to the initial state, a characteristic feature of the network termed perfect adaptation [13].

**Figure 1:**
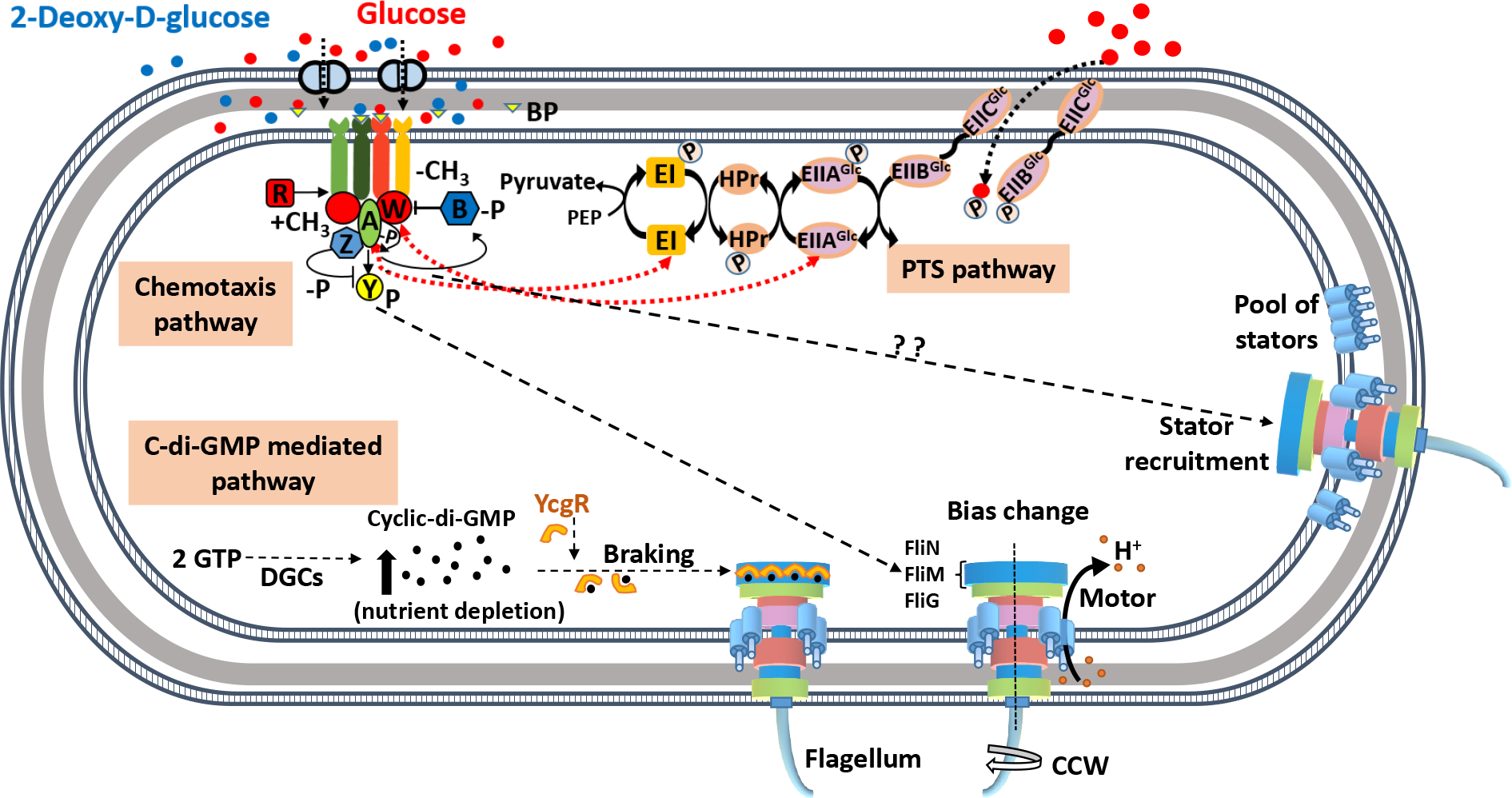
A schematic diagram of the known signalling pathways for control of locomotion in *Escherichia coli*. The chemotaxis pathway controls the direction of motor rotation via CheY protein which, on phosphorylation, binds to the FliM protein of motor-switch complex (FliG, FliM and FliN) and causes CW rotation of one or more flagella. The concentration of phosphorylated CheY (CheY-P) is dictated by the binding of ligands to their specific transmembrane receptors. Attractants such as glucose/2Dg bind to the periplasmic glucose-binding protein (BP) and are sensed by the Trg chemoreceptor protein. The binding results in a decrease in the autophosphorylation activity of CheA, which is part of CheA-CheW-receptor complex, leading to a lower concentration of CheY-P. The CheY-P binds to the FliM protein causing the motor to rotate in the CW direction. The proteins CheR and CheB control the degree of methylation of the receptors and allow the cell to adapt to the present concentration of attractant and sense subsequent changes. Additionally, glucose is a metabolizable ligand and is also sensed by an alternate pathway, namely, the phosphotransferase system (PTS), whose signals integrate into the chemotaxis pathway. The PTS generates a constant flow of phosphate groups via the PTS transporters (EI, HPr, and EII) from phosphoenolpyruvate (PEP) and activates the sensory complexes CheA-CheW thereby communicating with the motor via reduced levels of CheY-P. The CheZ protein accelerates the dephosphorylation of CheY-P. The motor rotation is generated via ion flow through the membrane-bound stators comprising of proteins, MotA and MotB while the speed is controlled via the intracellular concentration of the second messenger molecule, c-di-GMP. The latter is produced by diguanylate cyclase (DGC) enzymes from GTP and degraded by specific phosphodiesterases. Nutrient depletion conditions trigger production of c-di-GMP, which forms complex with YcgR protein and hinders the motor rotation. The present study suggests a signalling pathway that links the chemotaxis receptor to the motor speed via stator recruitment.

The torque of the flagellar motor is generated via ion flow through at least 11 membrane bound stators comprising of proteins, MotA and MotB [14, 15, 16]. Previous studies have demonstrated the discrete nature of torque generation by restoring the motility of paralysed cells (*mot* mutants) by induced expression of wild-type genes from plasmids. The latter caused a step-wise increase in motor speed, a process termed ‘resurrection’ [17, 18]. Recent experiments have shown that the number of bound stators vary with PMF with almost all stators leaving the motor following removal of PMF [19]. Consequently, the bacteria is able to vary the number of motor-bound stators depending on the external mechanical load [20, 21] although not all binding sites are necessarily occupied even at high load conditions due to the transient nature of stator binding. In addition to the torque variation with load, the speed of the motor is also controlled via the intracellular concentration of the second messenger Cyclic Diguanosine Monophosphate (c-di-GMP), whose concentration increases during nutrient depletion conditions [22, 23]. The c-di-GMP binds to the YcgR protein, a PilZ domain protein, which in turn interacts with the MotA to reduce the rotation speed of the motor. While the aforementioned studies have demonstrated the modulation of motor *bias* in response to sensing of ligand and change in motor speed with PMF or when exposed to starvation conditions, there is as yet no clear evidence of the chemosensory transduction system influencing the motor speed.

In this study, we investigate the role of sensing and metabolism of glucose on the motor performance both by measuring the motor speed and the concentration of stators bound to the motor via tethering experiments and confirming the same through run speed measurements in a population. For *E. coli*, glucose is the preferred carbon source for cell’s growth and there exist detailed studies on the molecular mechanism of sensing and adaptation to the ligand [24, 25, 26, 27, 28]. The signalling is achieved through two independent pathways, namely, through the phosphotransferase system (PTS) pathway and via the Trg-sensing pathway. The combined signals from the two pathways determines the phosphorylation state of CheA, which in turn determines the concentration of CheY-P, and therefore the motor bias. In order to differentiate the roles of sensing and metabolism on motor performance, we performed identical experiments with glucose and the non-metabolisable analogue of glucose, 2Dg. Adler et al [29] have shown that 2Dg is sensed by the Trg sensor, but unlike glucose, are not metabolized since cells do not grow in its presence. Further, cells lacking Trg show chemotactic response to 2Dg only when grown in the presence of glycerol through the activation of mannose PTS pathway [30].

The experiments were performed on the wild-type strain along with mutants lacking *trg* (Δ*trg*) and *ptsI* (Δ*ptsI*) genes. The influence of ligands on metabolism was assessed by measuring the membrane potential which corresponds to proton motive force in a population of cells. The data from the tethered cell experiments show a step increase in the motor speed when exposed to 2Dg. We also simultaneously measure the intensity of fluorescent dye-labelled stators bound to the motor (mutant strain JPA804) where increase in intensity indicates increase in number of stators. Our results indicate that mere sensing of a ligand can directly enhance the motor speed wherein the enhancement occurs due to an increase in the number of stators bound to the motor. These results clearly demonstrate the existence of a sophisticated signalling pathway that connects the chemotaxis signalling system to the torque generating stators.

## Results

### Sensing leads to increase in motor speed

In order to correlate the measured run speed with motor response, tethered cell experiments were performed. Figure 2A and 2B present the time evolution of the rotation speed after the introduction of glucose and 2Dg, respectively. In case of glucose, there is a sharp increase in rotation rate from an average speed of about 6 Hz to 7.5 Hz at about 100 s and again a further increase to about 9 Hz at around 650 s. In comparison, for 2Dg the rotation speed remains unchanged for about 300 s after which there is a sudden increase in average speed from about 4 Hz to about 5.7 Hz. The increased speed is sustained for about 200 s after which it returns to the pre-stimulus levels. The step-increases, which range between 1.0 to 2.0 Hz, are of magnitude similar to those observed in the resurrection experiments[17, 18].

**Figure 2:**
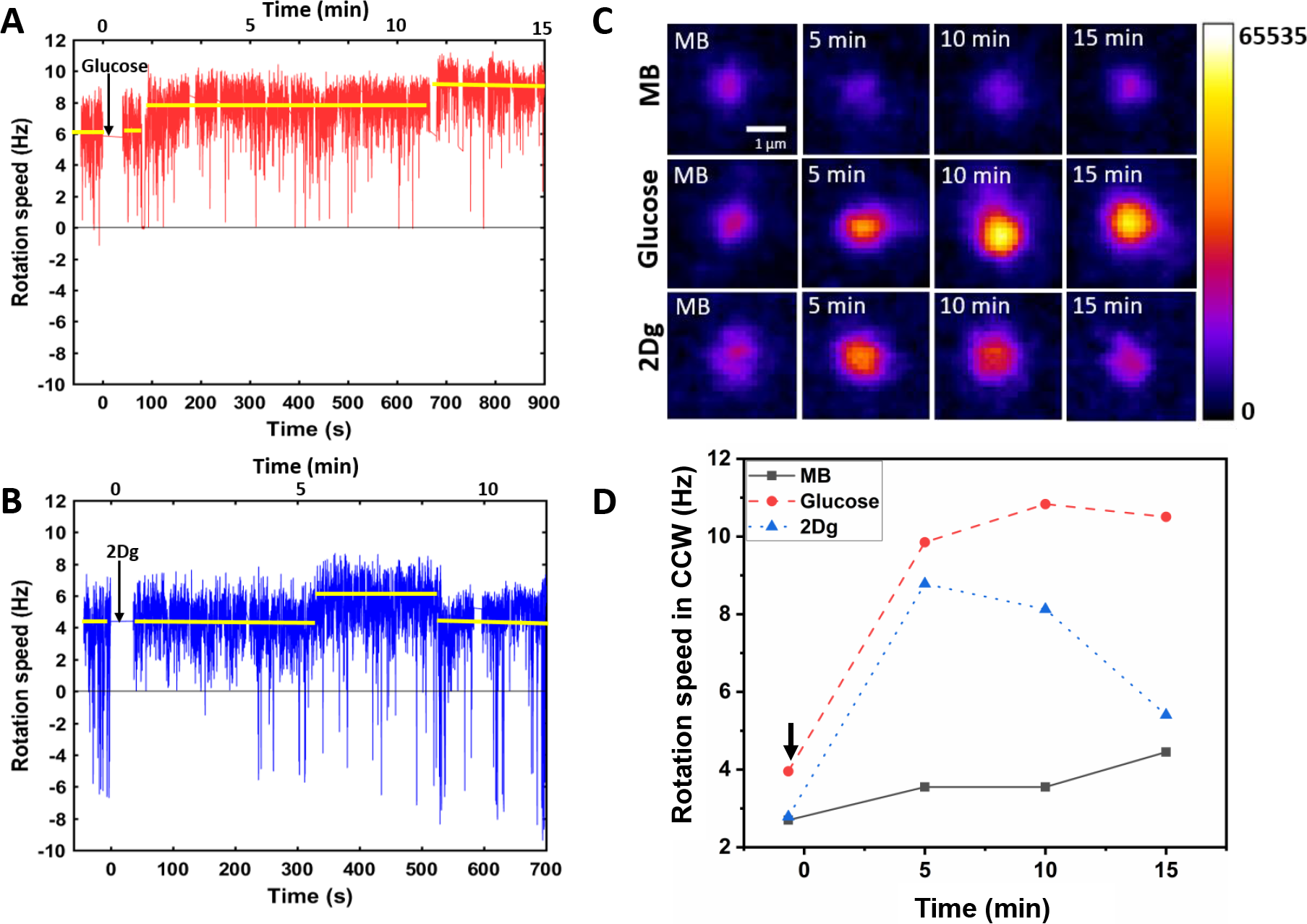
The motor rotation speed was measured after wild type (WT, RP437) cells were exposed (at *t* = 0) to 1000 *μ*M of (A) glucose, and (B) 2Dg. The arrows indicate the time of introduction of the ligands. The reported values of rotational rate are averaged over 0.16 s (10 frames). Imaging is performed about 50 s after the introduction of the ligand to let the accompanying flow to stop. (C) The fluorescence intensity of the stators (GFP-MotB in mutant strain JPA804) was measured at different time points after introduction of ligands and, (D) presents the corresponding motor rotation speeds (arrow indicates the time of introduction of ligand).

### Sensing leads to stator recruitment

The increase in the motor speed may be attributed to the recruitment of stator (MotAB) component of bacterial flagellar motor thereby increasing the torque on the motor. Therefore, GFP-tagged MotB component of stator for tethered mutant cell (JPA804) was directly visualised under a confocal microscope and the intensity variation at the motor site was recorded after the introduction of ligand in the medium (Figure 2C). While there is no change in the intensity for a cell in MB, the intensity increases with time when glucose is introduced. This observation is in agreement with previously reported experiments where the stator concentration increases with PMF[19]. Interestingly, on introduction of 2Dg, which is a non-metabolizable ligand, the intensity increases for a short duration (about 3-4 minutes) after which it falls to the pre-stimulus value. This result matches with the change in motor speed (Figure 2D) confirming that the modulation in motor speed is indeed caused by recruitment of stators upon mere sensing of ligand. Similar experiments were performed on about 7-10 motors and the relative average intensity values at fixed time points is plotted in Figure 3A. An increase of 23% at 5 min and 28% increase at 10 min was observed in the fluorescence intensity for glucose relative to its pre-stimulus value. An identical increase was observed at 5 min for 2Dg after which the intensity decreased to its pre-stimulus value. As expected there was no discernible trend in the intensity value measured for cell exposed to MB. Figure 3B presents the corresponding motor speeds for the cells, which show the same trend as that for the stator intensity. The normalized fluorescent intensity values in different motors for the three conditions and the corresponding normalized motor speeds are presented in Figure 3C. The intensity is proportional to motor speeds suggesting that stator elements are added following the introduction of the two ligands.

Following Samuel and Berg [31, 32], measurements of the variance in rotation speed of the tethered cells were also made as a function of mean number of revolutions to correlate response with number of active torque generating units. A large variance corresponds to few stator units powering the motor while a lower variance and therefore a smoother rotation implies larges number of stators. The measured variance followed the same trend as observed for the rotation speed and fluorescent stator intensity values in that the lowest variance was found in cells exposed to glucose followed by 2Dg and the highest in MB (Figure S9 C-E, Supplemental Information). The time evolution of the variance also matched with motor speed and stator intensity data.

**Figure 3:**
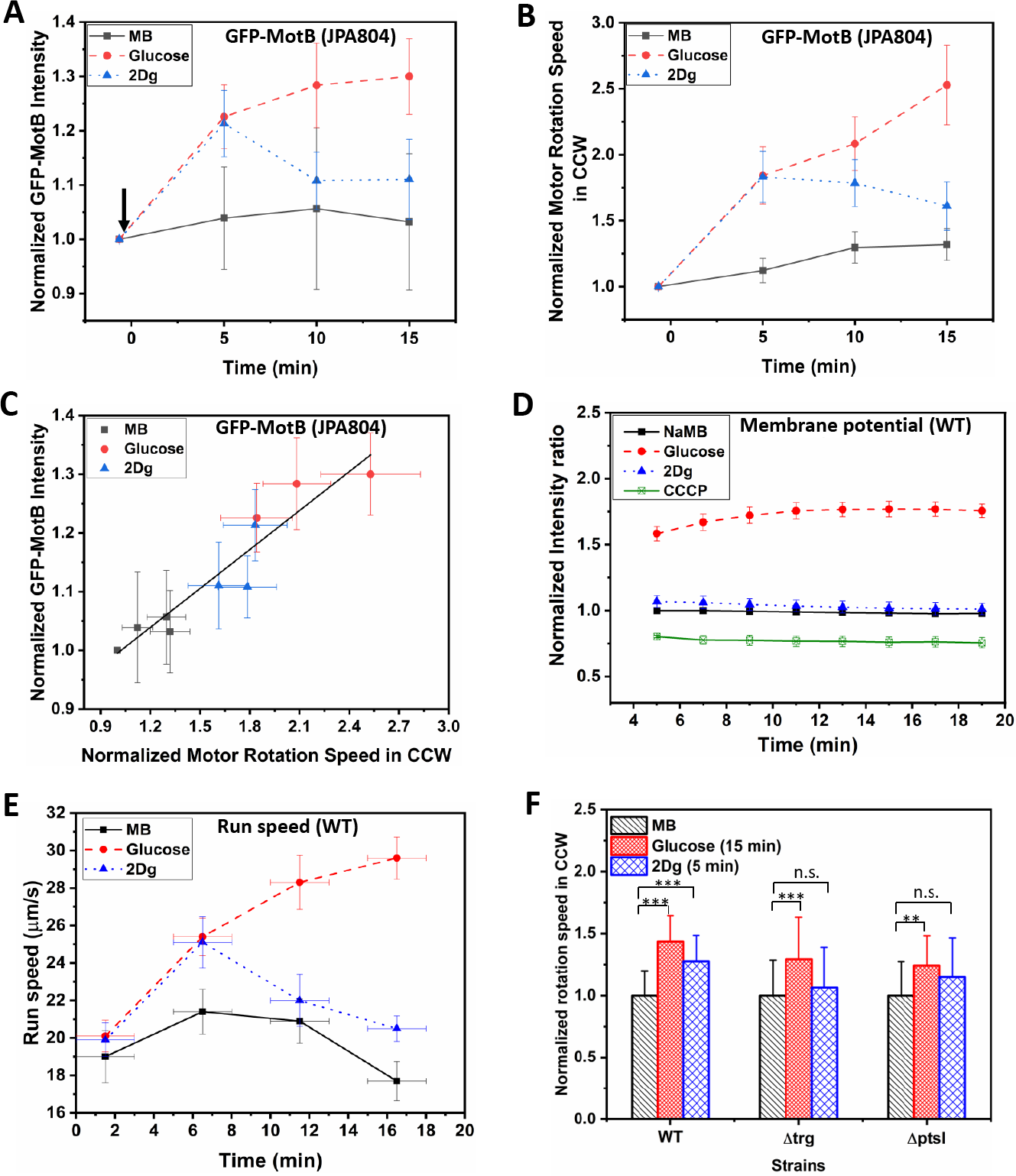
(A) Fluorescence (GFP-MotB) intensity variation in mutant strain JPA804 in 1000 *μ*M glucose, 1000 *μ*M 2Dg and motility buffer. Measured intensity of individual cells was first obtained in MB and then measured again for the same cells exposed to 1000 *μ*M of glucose and 1000 *μ*M 2Dg (arrow indicates the point of introduction of ligand). Increase in GFP intensity in glucose at all time points and in 2Dg at 5 mins are statistically significant at p < 0.01 computed by paired t-test. The error bars represent standard error of means from four independent experiments. The intensity of about 7-10 motors per ligand were measured. (B) The normalized average motor speed in CCW bias obtained from the tethered cell experiments presented in (A). (C) Motor intensity versus motor speed from data plotted in (A) and (B). The straight line is a linear fit. (D) Time dependent fluorescence intensity for membrane potential of WT in presence of 1000 *μ*M of glucose, 2Dg and sodium motility buffer for a population (3.75×10^8^ cells/ml). The error bars represent standard error of means from three independent experiments. Values are significant at p < 0.001 (one-way ANNOVA). (E) Measured run speed between consecutive tumbles for a population of WT cells in the presence of MB and when exposed to ligands. Each data point is obtained by averaging over 2500 cells. Speed increase in glucose for all three strains and the observed increase in 2Dg for the WT strain (at 5 min) as compared to MB is statistically significant at p < 0. 001. The Y-error bars represent standard error of means from four independent experiments. (F) Comparison of motor speed in CCW bias for the WT cell in the presence of ligands with those measured for mutant strains, Δtrg and Δ*ptsI* at fixed time points. Data represents value from at least 20 paired cells for each strain. Here, ‘***’ represents significance value p < 0.001 and ‘**’ represents p < 0.01 as measured by paired t-test. The highest speed for 2Dg were observed at 5 min.

### Sensing does not effect membrane potential

As discussed in the introduction, the stator recruitment has been directly correlated to PMF. To confirm that the observed increase in the presence of 2Dg was not due to changes in PMF, we measured the change in transmembrane electrical potential and pH gradient (Δ*pH*) in the presence of glucose and 2Dg. Since *E. coli* is a neutraphile, the change in the PMF is primarily due to changes in Δ*ψ* (Figure S4, Supplemental Information) [33]. Figure 3D presents fluorescence intensity values for WT, which are proportional to membrane potential. The buffer solution in these experiments contain sodium salts (NaMB) instead of potassium since the latter has been shown to interfere with membrane potential measurements[34, 35]. The measured potential is the lowest for the control, namely, CCCP (Carbonyl cyanide m-chlorophenyl hydrazone), which is an ionophore and eliminates the membrane potential. While there is no change in the potential for NaMB and 2Dg and their values are very close, the intensity value in presence of glucose is significantly higher. This clearly demonstrates that the stator recruitment in presence of 2Dg is independent of PMF and is caused solely by sensing of 2Dg. For glucose, the increase in motor speed would be a combination of PMF and that due to sensing.

### Average swimming speed in a population

Given the above observations, does the single motor behavior influence the swimming speed cells? In order to address this question, run speeds were measured at constant ligand concentration at four different time points, namely 1.5, 6.5, 11.5, and 16.5 min after the cells were exposed to ligands (average of over 2500 cells). In presence of glucose, the run speed increased monotonically (Fig 3E) with almost a 50% increase (29.6±1.0*μ*m/s, mean ± s.e.) at 16.5 min compared to that measured at the start of the experiment (20.1±0.9*μ*m/s). In case of 2Dg, the run speed increased upto 6.5 min reaching a value identical to that observed for glucose at the same time, resulting in a 26% increase (25 ±1.4*μ*m/s) compared to the speeds measured at the start of the experiment. However, at longer times, unlike in glucose, the run speed decreased and reached values observed at the start of the experiment. In the absence of ligands, the run speed showed negligible change with speed ranging approximately between 18 to 21 *μ*m/s for the entire duration of the experiment. These results indicate that sensing alone can enhance the run speed, albeit for a short duration, as observed in case of 2Dg. A further increase in run speed is observed in case of glucose suggesting that metabolism plays a role in sustaining a high run speed. Clearly, the observations made on a single motor via tethered cell experiments indeed reflect in the population behavior wherein stator recruitment due to sensing alone can enhance the swimming speed of a population. We note in passing that the concentrations of the ligands are sufficiently low that osmotic effects do not influence the motion [36].

### Role of Trg receptor and PTS in speed modulation

In order to characterise the role of Trg receptor, a mutant strain lacking in *trg* was exposed to both glucose and 2Dg and motor speed was recorded at various times (Fig 3F). There was no difference in response between MB and 2Dg for the Δ*trg* mutant strain. However, in glucose, there was significant increase in the motor speed (about 30%) compared to the pre-stimulus value. This is in contrast to that observed in wild type where the motor speed increased in both glucose (about 40%) and 2Dg (about 30%). These results clearly show that in the absence of the Trg sensor, the motor speed remains unaffected in the presence of 2Dg thereby reconfirming the earlier observation that sensing alone can contribute to motor speed modulation. Experiments measuring the membrane potential show a behavior similar to that observed for wild type cells (Fig 4A). This is expected since 2Dg is not metabolized and hence does not contribute to PMF and consequently, presence or absence of Trg sensor has no effect in presence of 2Dg. The mutant strain demonstrates increased PMF in glucose through metabolism although the ligand is not sensed. This behavior is observed at the population level (Fig 4B), wherein there is no change in run speed in response to 2Dg, which is identical to that observed in MB, while there is a significant increase in the presence of glucose (about 50% increase).

**Figure 4:**
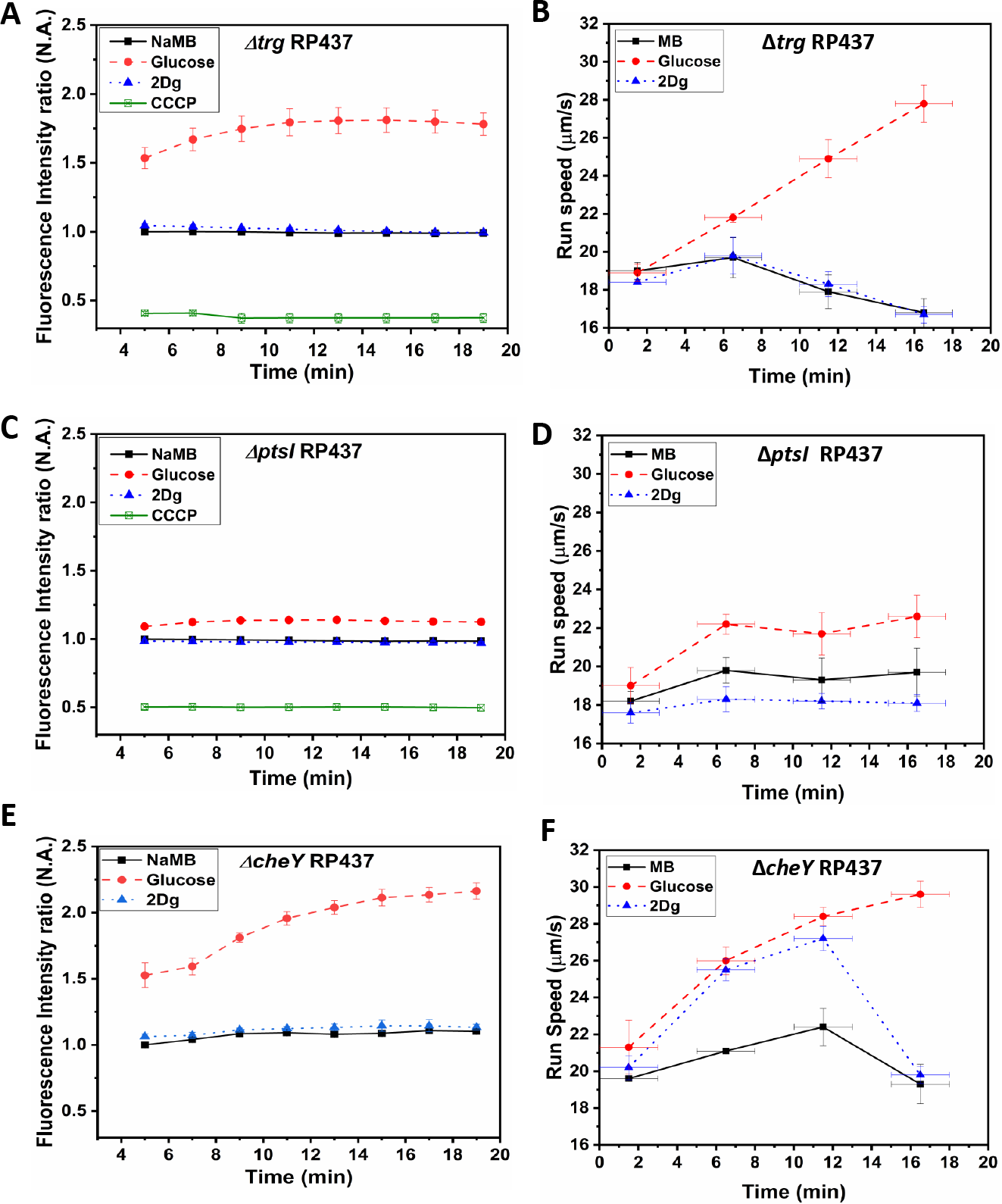
Time dependent fluorescence intensity for membrane potential and time dependent run speed variation for mutant strains, (A) and (B) Δ*trg* RP437, (C) and (D) Δ*ptsI* RP437, and (E) and (F) Δ*cheY* RP437 in presence of MB, and 1000 *μ*M of glucose and 2Dg. Final concentration of cells for membrane potential measurement was 3.75 × 10^8^ cells/ml and the values for glucose compared to that for MB are significant at p < 0.001 (one-way ANNOVA). Each data point in the run speed plot is obtained by averaging over 2500 cells. Speed increase in glucose for all three strains and the observed increase in 2Dg for the WT strain and CheY mutant (at 5 min) as compared to MB is statistically significant at p < 0.001 (one-way ANNOVA). The Y-error bars represent standard error of means from four independent experiments, while the X-error bars represent the duration of 3 minutes over which the measurements were made for each data point.

In order to estimate the role of metabolism in motor speed modulation, we performed the same set of experiments in mutants lacking the *ptsI* gene. There was a marginal increase in motor speed (Fig 3F) and run speed (Fig 4D) in presence of glucose while no significant change was observed in the presence of 2Dg. It can be noted that the membrane potential dropped drastically due to the absence of *ptsI* gene (Fig 4C) compared to that observed in either wild type or Δtrg strain. The small increase in membrane potential measured for the mutant strain in the presence of glucose explains the marginal increase in the motor and run speed.

### Role of CheY

To determine whether CheY is necessary for the observed increase in run speed (Fig 3E), experiments were also performed with a strain lacking CheY (Δ*cheY* mutant). These mutant cells do not tumble but always run since the flagellar motors rotate in the CCW direction in absence of CheY-P. The results of such experiments are shown in Fig 4F. The trends are very similar to those observed with WT (Fig 3D) in that the run speed increases monotonically with time in presence of glucose while the speed first increases and then decreases in case of 2Dg. The similarity of the observed trend with WT indicates that CheY-P has no role in the observed modulation in motor speed. The trend in membrane potential for ΔcheY in presence of both ligands was similar to that observed in the wild type strain (Fig 4E).

## Discussion

It is well known that *Escherichia coli* modulates its motor bias in response to ligands [1]. For example, in case of glucose, which is sensed via the Trg chemoreceptor, sensing leads to the dephosphorylation of CheY-P which in turn reduces the motor reversals leading to smooth runs. Glucose is also sensed and consumed via the PTS pathway, which is known to regulate CheY-P, thereby altering the bias. Neumann et al [28] have shown that the signals from the Trg receptor and the PTS pathway combine (additive) to determine the reversal frequency.

Although it is well known that the chemotactic pathway modulates the motor bias, the existence of a pathway that also modulates the motor speed is not known. Previous studies have shown that in tethered cell experiments, where the torque is in the high load regime, the motor speed is proportional to the ion motive force in both proton and sodium-driven motors [7, 37]. These changes in motor output have been attributed to the change in the number of stators, which are assembled from a mobile pool of MotB molecules over a time scale of minutes when the PMF is increased [15, 38]. In contrast, for cases where the mechanical load is low, such as in bead-assay with sub-micron sized beads, the motor output can also vary with the magnitude of the load due to load-dependent stator recruiment [20, 21]. Thus this dynamic engagement of stators has been shown to modulate motor speed in response to both nutritional status and mechanical load although the exact mechanism responsible for them are not known. While these studies clearly indicate the role of metabolism and mechanical load in motor speed modulation via stator recruitment, there is no evidence hitherto that mere sensing of a ligand can directly influence the motor speed at a fixed mechanical load. The tethered cell experiments reported here clearly demonstrate the recruitment of stators in response to sensing of a ligand and therefore suggest the existence of a pathway that links the sensor to the motor.

Our experiments show an increase in stator density at the motor in response to 2Dg, a non-metabolisable analogue of glucose. This increase in stator density led to a concomitant enhancement in motor speed and thereby the run speed in a population. Since 2Dg is sensed by the Trg sensor, this indicates that a signal for speed modulation is communicated to the motor from the sensor. Further, there was no change in membrane potential confirming the absence of role of metabolism and thereby suggesting that any change in the motor speed is only due to the sensing mechanism. This was further confirmed by deleting the *trg* gene, which eliminated changes in motor speed in presence of 2Dg. In case of glucose, the enhancement in motor speed is much greater compared to that in 2Dg due to the combined effect of sensing and metabolism, caused by an even greater recruitment of stators (as indicated by increased intensity of fluorescently tagged stator). Here, the increase in motor speed is due to glucose metabolism leading to increased PMF [39] while that due to sensing is through the Trg receptor. When the metabolism was disrupted (Δ*ptsI* strain), the basal values of the motor speed reduced significantly. A small increase in motor speed was observed when the cell was exposed to glucose but changes were statistically insignificant in the presence of 2Dg. The weak response observed for the Δ*ptsI* strain may be due to the impaired chemotaxis pathway since PTS genes are known to control the expression of proteins involved in chemotaxis [28].

An alternate method to characterise stator recruitment is to determine the variance in mean rotation rate of the motor as a function of the number of revolutions[31, 32]. Thus, a smaller variance would indicate an increased number of stators, thereby making the motor rotation smoother. The tethered cell experiment with the wild type cells show clearly a decrease in variance when exposed to 2Dg (Fig S9 B and E) compared to that in MB. The variance decreased even further when exposed to glucose indicating a higher recruitment of stators due to the combined effect of metabolism and sensing (Fig S9 A and D). These observations are consistent with the stator density measured via fluorescence.

Could the observed modulation of motor speed be controlled by the well-known chemotaxis pathway where the phosphorylated CheY binds to the motor to modulate not only motor bias but also motor speed? To answer this question, motor speed of a mutant lacking in cheY was measured in the presence of both 2Dg and glucose. The observed motor speed modulation was identical to that for the wild type indicating that CheY does not play role in stator recruitment. These observations clearly suggest that there exists a signalling pathway, independent of the chemotaxis pathway, that links the sensor to the motor. Note that in the cheY mutant, the bias is locked in the CCW direction and therefore is also always in the run mode. Experiments with wild-type cells on the other hand show a momentary increase in CCW bias after which the bias recovers to the pre-stimulus value thereby demonstrating the well known chemotactic response. This was observed for both glucose and 2Dg albeit a larger increase in CCW bias in the former (see supplementary information, Fig S6).

The above results may be understood by considering the state of the motor on the torque-speed curve (Fig 5A). The dashed line originating from the origin is the load path of the motor under high load conditions. When the cell is exposed to 2Dg, one or more stators are recruited temporarily (state 1 to 2 in Fig 5A, blue line) thereby increasing the rotation speed. The stators disengage within a short time (about 5-10 min) after which the rotation rates return to the pre-stimulus values (back to state 1), see schematic in Figure 5B. This effect is due to sensing alone. However, when the cell is exposed to glucose, a metabolite, the combined contribution from sensing and PMF leads to a much larger increase in the number of bound stators (state 1 to 3, red line) and the speeds remain high for a much longer time periods (over 15 min), due to the metabolism.

**Figure 5:**
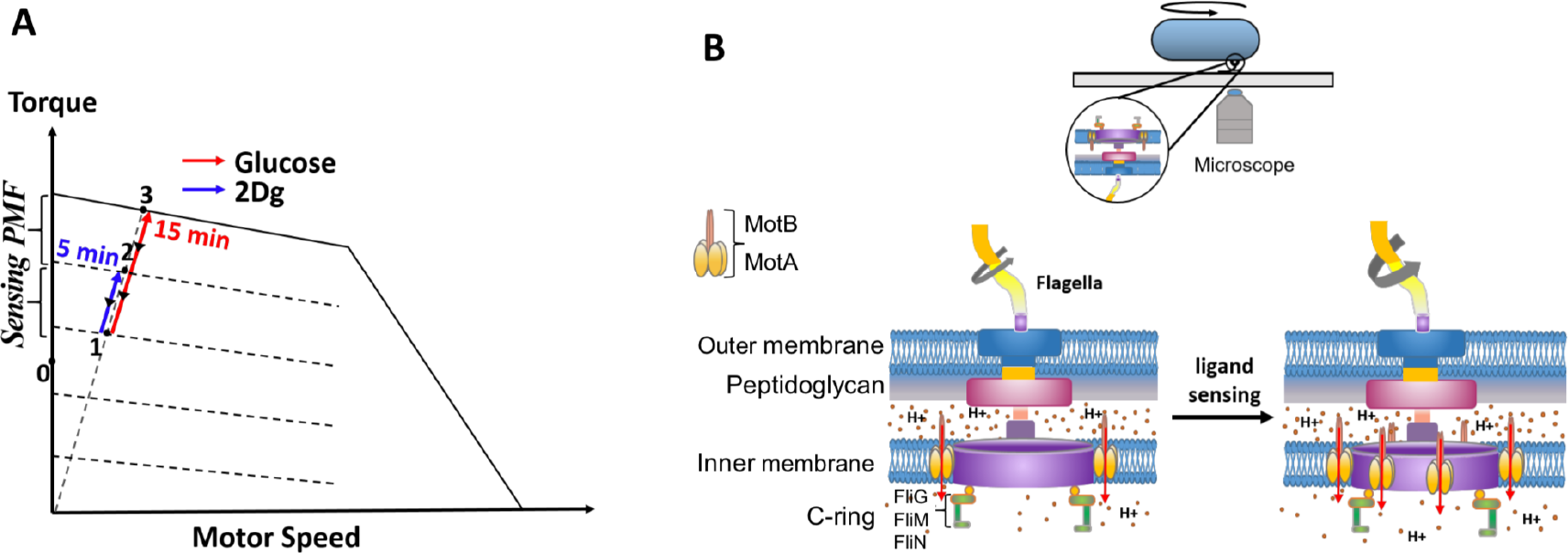
(A) A representative plot of steady-state torque-speed curves for CCW rotation of the bacterial flagellar motor at varying loads along with the expected state of the motor along the load line at high load when the cell is exposed to 2Dg and glucose. (B) A schematic of the bacterial motor along with a cell tethered to a glass coverslip is included to the right. Motor speed increases as a result of stator recruitment upon ligand sensing.

At the population level, these results imply a more efficient chemotaxis response since the drift velocity of a population of cells in response to a ligand gradient is dependent on both the tumble frequency and the swimming speed[40]. Further, the pathway is independent of the known sensory transduction pathway since the observed speed modulation also occurs in the absence of CheY (Fig 4F). Recent studies have shown that c-di-GMP effector YcgR can inhibit flagellar motility by interacting directly with the motor to modulate both its bias and speed [22, 23, 41]. While the aforementioned mechanism has been shown to inhibit motor speeds there is no evidence that the same can enhance motor speeds beyond pre-stimulus levels. Irrespective of the details of the mechanism, the results presented here point to a sophisticated signalling network with multiple strategies to regulate motor behavior enabling an efficient response to changes in bacteria’s environment.

## Materials and Methods

### Bacterial strains and growth condition

*E. coli* RP437 and its four mutants (see supplemental information, Table S1) were inoculated from glycerol stock into tryptone agar plates. A single colony isolate was grown overnight and then sub-cultured in 50 ml tryptone media (0.015g/l; Sigma) till mid-log phase (OD _600_ at 200 rpm, 30°C).

### Tethered cell experiment

Cells were centrifuged at 4000 rpm and washed twice with motility buffer (MB). Cells were then resuspended in 1 ml MB and passed 75 times through a syringe fitted with 21-gauge needle to enable shearing of flagella. Cells were centrifuged at 4000 rpm for 10 min to obtain cells having short flagellar stub. The cell pellet so obtained was again diluted in MB.

Rectangular chambers were created with a microscope cover slip as base and flexible thin sheet of polydimethylsiloxane (PDMS)(Sylgard 184; Dow corning) as the top wall separated by two parallel silicon rubber spacers of approximate thickness 400 *μ*m. The thin sheets of PDMS were prepared by spreading the uncured polymer on polystyrene petri plates and then curing them for 8 h at 50 °C. The microscopic coverslip was cleaned with ethanol and smudged with thin layer of silicone oil [42]. Use of silicone oil prior to antibody treatment in our case resulted in a more stable interaction between anti-flagellin antibody and the glass slide. This ensured that tethered cells were not washed away when the ligands were introduced.

The aforementioned arrangement results in a chamber that is permeable to oxygen. The working volume was of the chamber was approximately 80-100 *μ*L. The chamber was first incubated for 30 min with a solution containing anti-flagellin antibody (Abcam) specific for bacterial flagellin. The solution was prepared by diluting the antibody 1000 folds in 50mM Tris-HCl (pH 7.6). The chamber was washed gently with MB after which sheared cells were introduced into the chamber. An incubation time of 15-20 min was sufficient for the flagellum to tether to the antibody-coated coverslip surface. Unattached cells were washed and the channel was filled again with MB. Once the chamber was filled with MB, concentrated ligand solution (5000 *μ*M glucose or 2Dg) of a known volume was then introduced gently through four holes punched in the PDMS sheet so that the average concentration of the final solution in the chamber is 1000 *μ*M. The four holes surrounded the visualization region and were about 3-4 mm from the region. Experiments with color dyes showed that a combination of convective flow induced by the incoming dye solution and accompanying diffusion led to the dye reaching the visualization region within 30-45 seconds of introducing the dye. In order to measure the response of single flagellar motor to ligands, we first measured the response in MB and again after adding glucose/2Dg. Such pair-wise assessment removed any variability arising due to cell size and tethering geometry.

Experiments with ligands were performed at an ambient temperature of 23 ° ± 1C. The imaging was done with an inverted microscope (Olympus IX71) fitted with 100X (1.4 NA), oil-immersion objective. Area was scanned for rotating cells and videos were captured at an acquisition rate of 60 fps for 30 to 44 s using a CMOS camera (Hamamatsu Inc).

### GFP-MotB intensity measurement

For GFP-MotB intensity measurement, cell preparation was done in the same manner as that for the tethered cell experiment except that antiflagellin antibody was not used in the flow chamber. This because the mutant strain JPA804 (RP437 MotB-GFP tagged cell) has sticky flagella [38]. Cells were imaged via spinning disc confocal microscopy (Perkin Elmer UltraVIEW system with Olympus IX71) under 100X (NA 1.4) oil immersion objective. The cells were excited with a 488 nm-laser with exposure time of 200ms (4-5 mWatt). Images of 16-bit resolution were captured at an acquisition rate of 6 fps for 1 second at different time points for rotating cells using an EMCCD camera (Hamamatsu Inc). Simultaneously, bright field videos were captured using a CMOS camera (Hamamatsu Inc) at an acquisition speed of 34 fps for 5 seconds for measuring rotation speed. Videos were analysed using ImageJ software and the fluorescence intensity of the stator proteins and motor speed were determined for the rotating cells. All the experiments were performed at 25 °C. To determine the fluorescent intensity of the motor at a given time, we determined the highest intensity pixel in the motor region and subtracted the background intensity obtained from the region surrounding the bacterium. We observed that in some cases the cells did not rotate for the entire duration of the experiment. Such cases were not included. For analysis, we chose cells that were tethered at their polar ends and ignored cells that were tethered at the center of the body.

### Determination of swimming speed

Cells were centrifuged at 4000 rpm and washed twice with motility buffer (MB). Cells were added to each vial of ligand (1000 *μ*M of glucose and 2Dg) and MB such that the cell density was low, about 10^6^-10^7^ cells/mL. The cells were introduced into glass, rectangular micro channels (5 cm (L) × 1000 *μ*m (W) × 100 *μ*m (H)) via capillary action [43] at different time points (t =0 to 15 min) for visualization and measurements. Both ends of microchannel were sealed with wax after introduction of cells. Trajectories of swimming bacteria were recorded with 40X (0.75 NA) objectives using dark field microscopy. At each time point, six videos of about 15 s duration were recorded over a span of 3 min at a frame rate of 21 fps. Images were taken in the central region of the micro channel away from the channel walls. The trajectories of the cells were obtained using a commercial software, ImageProPlus. The data was analysed using an in-house code written in MATLAB to obtain the swimming speed and cell orientation from more than 2,500 cells for each condition. We considered only those cells that were within 1*μ*m of the focal plane with tracks longer than 0.5 s, which ensured that all out-of-plane motions were ignored by the analysis. A tumble event was identified when the swimming speed of the cell was below half the mean swimming speed and the change in the turn angle was greater than *4°* between successive frames (at 21 fps)[44]. The measured average swimming speed of 18.2±7.9*μ*m/s (average ± standard deviation) and an average turn angle of 71° for RP437 cells dispersed uniformly in a microchannel containing plain motility buffer are close to those observed for the same strain reported earlier, 18.8±8.2*μ*m/s and 69°, by Saragosti et al.[45]. The details of the analysis has been described in earlier published reports [43, 46].

### Measurement of membrane potential

Cells preparation method was similar to that described before except that sodium phosphate buffer (NaMB) was used instead of the standard MB. NaMB contained Na_2_HPO_4_, 1.64 g; NaH_2_PO_4_, 0.47 g; NaCl, 8.77 g and EDTA, 0.029 g per litre of distilled water. The chemicals Na_2_HPO_4_, NaH_2_PO_4_ and NaCl were obtained from Merck^®^ while EDTA and Carbonyl cyanide m-chlorophenyl hydrazone (CCCP) were purchased from Sigma Aldrich^®^. The cell pellet was dissolved in 500 *μ*l of NaMB and 3.75 *μ*l of the resulting solution was further diluted in NaMB containing ligand at a 1000 *μ*M concentration such that a final cell density of 3.75×10^8^ cells/ml was achieved.

3,3-Diethyloxacarbocyanine iodide (DiOC_2_(3)) from ThermoFisher Scientific^®^) is a lipophilic cationic dye and was used as a membrane potential probe. DiOC_2_(3) was added at a final concentration of 30 *μ*M to each sample vial after which samples were taken for fluorescence readings in microplate reader (Varioskan LUX plate reader) with excitation at 488 nm. Further, addition of EDTA facilitates dye uptake in gram negative cells. Hence, 100*μ*M EDTA was introduced in sodium phosphate buffer.

DiOC_2_(3) enters the cell as a monomer and binds to the membrane. Upon excitation by 488nm wavelength light, the dye emits both green (520nm) and red (620nm) fluorescence. The green fluorescence intensity is dependent on the size of the cell while the red fluorescence intensity is dependent on both the membrane potential and the size of the cell. The ratio of red and green fluorescence (R/G) eliminates the dependence on cell size and directly correlates with the membrane potential of the cells. Membrane potential is negative on the inside of the membrane and so the magnitude of the negative charge correlates with the magnitude of R/G[47, 48, 49]. CCCP is an ionophore which increases membrane permeability to protons and thereby eliminates membrane potential. In such a case, the R/G is the lowest and is used as a control in our experiments.

## Acknowledgement

Financial support from the Department of Science and Technology, India (SB/S3/CE/089/2013) and Department of Biotechnology, India (BT/PR7712/BRB/10/1229/2013) is acknowledged.

## Notes

### Competing Interest Statement

The authors have declared no competing interest.

### Summary of Updates

In the revised manuscript we have included a new Fig 1, which summarises our current understanding of the various pathways responsible for locomotion. An additional subplot (fig 3C) is included where the intensity vs motor speed is plotted for all three cases (glucose, 2Dg and MB). The points lie on a straight line suggesting that stator recruitment is responsible for motor speed increase in all three cases.

