## Supplemental Information for "Ligand Sensing Enhances Bacterial Flagellar Motor Output via Stator Recruitment"

### Materials and Methods

*a. Construction of gene knockout strains* All mutants are derivatives of *Escherichia coli* RP437 strain as described in Table S1. The $\Delta cheY$ mutant was gifted by Professor J. S Parkinson and JPA804 strain by Professor Judith Armitage. Other knockout mutant strains were developed by using standard site-specific recombination method for inactivation of genes [1, 2]. The desired gene was deleted by replacing it with kanamycin marker. The kanamycin marker was further removed by site specific recombination around FRT region leaving behind scar of 80 base pairs, using a plasmid expressing an FLP recombinase. The desired deletion was confirmed by colony PCR with oligonucleotides mentioned in Table S2. The details of the PCR with verification primers is given in Figure S1.

TABLE I. *E. coli* strains used in this study

| Strain | Relevant genotype | Parent strain | Comments, references |
|---|---|---|---|
| RP437 | | | Wild type [3] |
| RP5232 | $\Delta cheY$ | RP437 | [3] |
| JPA804 | GFP MotB FliC-sticky | RP437 | [4] |
| MTKV01 | $\Delta trg$::Frt | RP437 | This study |
| MTKV04 | $\Delta ptsI$::Frt | RP437 | This study |

TABLE II. Oligonucleotides used as primers for gene knockout

| **Designation** | **5'– 3' sequence** | **Function** |
|---|---|---|
| **PtsI- F1** | TAATTTCCCGGGTTCTTTTAAAAATCAGTCACAAGTAAGGTAGGGTTATG ATTCCGGGGATCCGTCGACC | *ptsI* deletion |
| **PtsI-R2** | AAGCAGTAAATTGGGCCGCATCTCGTGGATTAGCAGATTGTTTTTTCTTC TGTAGGCTGGAGCTGCTTCG | |
| **Trg-F1** | GCCGATGACTTTCTATCAGGAGTAAACCTGGACGAGAGACAACGGTAATGATTCCGGGGATCCGTCGACC | *trg* deletion |
| **Trg-R2** | GGGATCTGTCGATCCCTCCTTGAACATTTTCACACCGTAGCGAAACTAACTGTAGGCTGGAGCTGCTTCG | |
| **PtsI- VF1** | CTGCTGCCCAGTTTGTAAAA | Confirming *ptsI* deletion |
| **PtsI- VR2** | TTTACCAATGGTGCCGTCTA | |
| **Trg-VF1** | AGGCATCCTATGAGGTTTCCT | Confirming *trg* deletion |
| **Trg-VR2** | CTATCTCGTCAACTTACGGTTGAAT | |

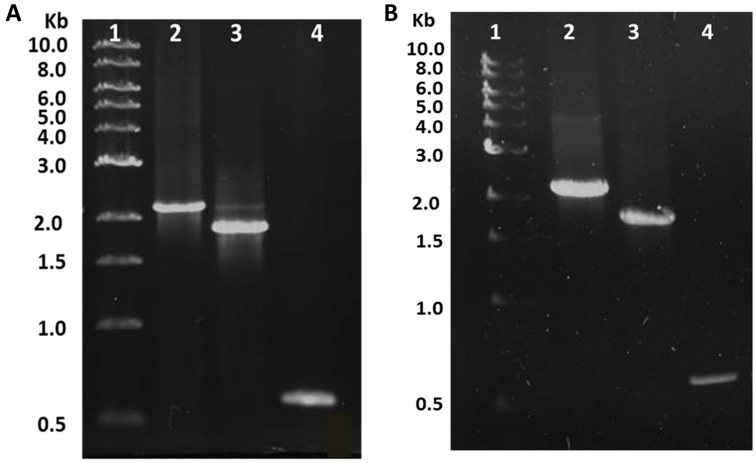


FIG. S1. Agarose gel (0.8%) showing the result of PCR with verification primers. (A) Mutation of *trg* gene in *E.coli*, Lane 1. Marker, 2. RP437 wildtype (WT) with *trg* gene (2.1 kb), 3. Δ*trg*::kan (1.9 kb), 4. Δ*trg* (600 bp) (B) Mutation of *ptsI* gene in *E.coli*, Lane 1. Marker, 2. WT with *ptsI* gene (2.2 kb), 3. Δ*ptsI* ::kan (1.9 kb), 4. Δ*ptsI* (600 bp)

*b. Tethered Cell Experiment* Figure S2A shows a schematic of the top and side view of a cell tethered to a glass coverslip [5]. Figure S2B presents an image of the cell along with two circles drawn by the Particle Tracker plugin of ImageJ software for identifying the cell ends. The rotation of the cell was recorded and the angular position, obtained as a function of time, was used to determine the rotation speed and the rotation bias.

*c. pH homeostatsis* WT RP437 cells transformed with plasmid pMS201 having a gene with kanamycin resistance and green fluorescence protein (GFPmut2) was gifted by Professor Supreet Saini. The strain was used for determining pH homeostasis. Experiments were performed in presence and absence of 20 mM sodium benzoate (weak permeant acid),

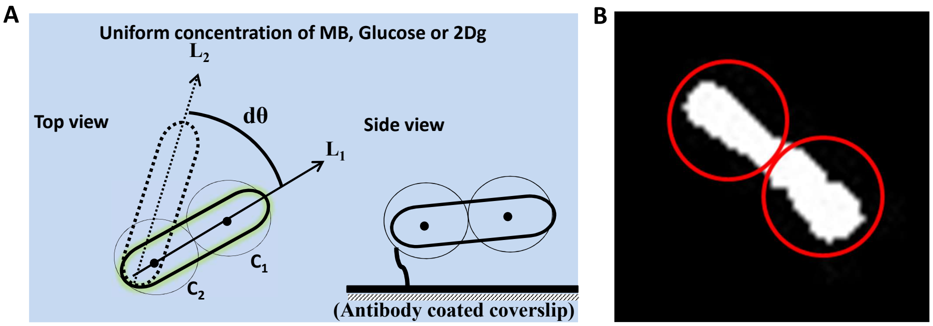


FIG. S2. (A) Illustration for motor speed calculation for a tethered cell (top and side view). Cells can be either tethered at the center or at the pole of the cell to an antibody-treated glass surface. In the acquired images, two circles are fitted along the length of the cell and their centers, $C_1$ and $C_2$, are tracked in each frame. The line joining the centers is tracked in time and is used to determine the rotation speed and the direction of rotation (between lines $L_1$ and $L_2$). (B) Top view of the cell fitted with two circles using Particle Tracker plugin for ImageJ software.

which is known to collapse the pH difference across the cell membrane and equilibrate it to the pH value of the external medium. The cells were suspended in motility buffers with different pH ranging from 5.5 to 8.5. Samples were excited at 485 nm and emission was collected from 495 nm onwards using a fluorescence spectrophotometer (Varioskan LUX plate reader). Peak emission was observed at 508 nm and the same was used for analysis.

### Image Analysis

*d. Motor speed measurement* Several videos of rotating cells were captured for analysis. Individual cell was cropped from the main movie and the resulting movies were thresholded in ImageJ (NIH). The ImageJ plugin, Particle tracker 2D/3D, was used to track the rotation of cell in each frame [6]. Particle detection and linking parameters were adjusted in such a way that only two circles fit the two ends of the rod shaped bacteria (Fig S2A, C). The coordinates of the centers of the two circles gave the orientation angle of the cell (Fig S2A). The change in orientation angle was determined for each subsequent frame (d$\theta$) and was used to determine the direction of rotation and the instantaneous rotation rate (Hz). Rotation rate values between -0.05 and 0.05 Hz were considered as a pause [7]. Moving average of 10

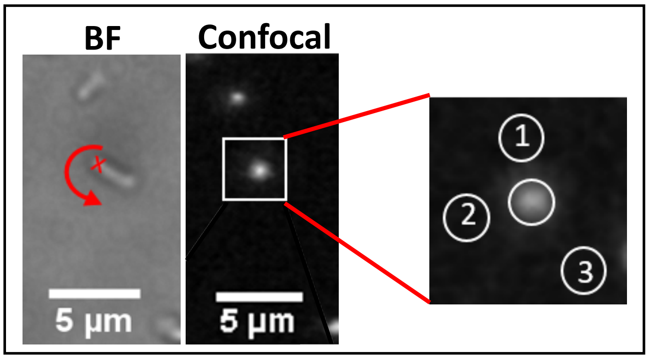


FIG. S3. The left panel corresponds to the brightfield image while the middle panel is the confocal image of the same location. The red cross indicates the location of the motor while the arrow indicates the direction of rotation. The fluorescence intensity of GFP-tagged MotB motor is obtained by noting the maximum intensity value in the motor region and then subtracting the average pixel intensity value from it. The latter was obtained from three, evenly spaced, neighboring circular patches around the cell, as shown in the enlarged image of the cell of interest (right panel). The average intensity value was independent of the exact location of the patches as long as they were evenly spaced.

instantaneous speeds were calculated so as to obtain the mean rotational speed and CCW bias. Since a cell undergoes several revolutions during the course of the experiment, the mean time required to complete a specified number of revolutions and the corresponding variance was determined from image analysis.

*e. GFP-MotB intensity analysis* Confocal microscope was used to capture 6 to 7 images at 6fps at each time point. Next, the maximum intensity in the motor region was determined from the image stack. To eliminate the effect of varying background intensity from one experiment to another, the average pixel value obtained from three distinct locations (highlighted by the circles in Fig S3) around the cell was subtracted from the image. This ensured that the fluorescence intensity of the stators was not influenced by the intensity variation in the background. This process was applied to all cells. The same cell was also visualized in brightfield to simultaneously capture the motor rotation speed.

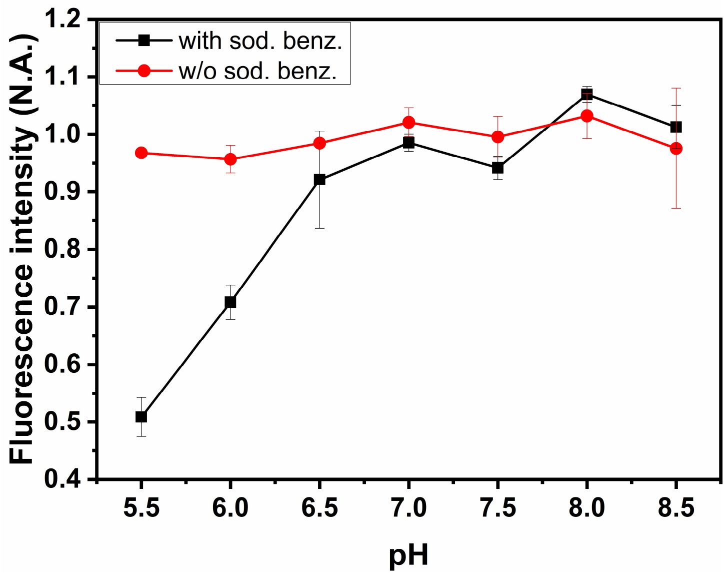


FIG. S4. Response of RP437 cells to motility buffers with different pH values when exposed to sodium benzoate. Cells exhibit homeostasis when suspended in different pH buffers in absence of sodium benzoate. Each data point corresponds to cells at OD 0.3. The error bars represent standard error of means from three independent experiments.

**Additional Results**

*f. pH homeostasis in E. coli* Figure S4 presents the fluorescent intensity corresponding to the cytoplasmic pH. In the absence of sodium benzoate, the intensity value is independent of the external pH indicating that the cell maintains a constant pH irrespective of the external conditions. In the presence of sodium benzoate, which causes the pH difference to collapse, the fluorescence intensity varies with change in external pH. These observations are in agreement with previously published results [8–10]. As a consequence, all changes in the PMF observed in our experiments, which were all performed near neutral pH, are due to changes in membrane potential.

*g. Motor speed of WT in glucose after 5 min of incubation* Figure S5 presents the rotation rate and the CCW bias for WT cells before and after they were exposed to glucose. The measurements were made 5 min after exposure to glucose. As expected, both the rotation rate and CCW bias are higher compared to the pre-stimulus value. The corresponding measurements for 2Dg have been reported in the main text (Fig S7C) and (Fig S8C). Note that rotation rate for glucose at 15 min (Fig S7B) is higher than that at 5 min.

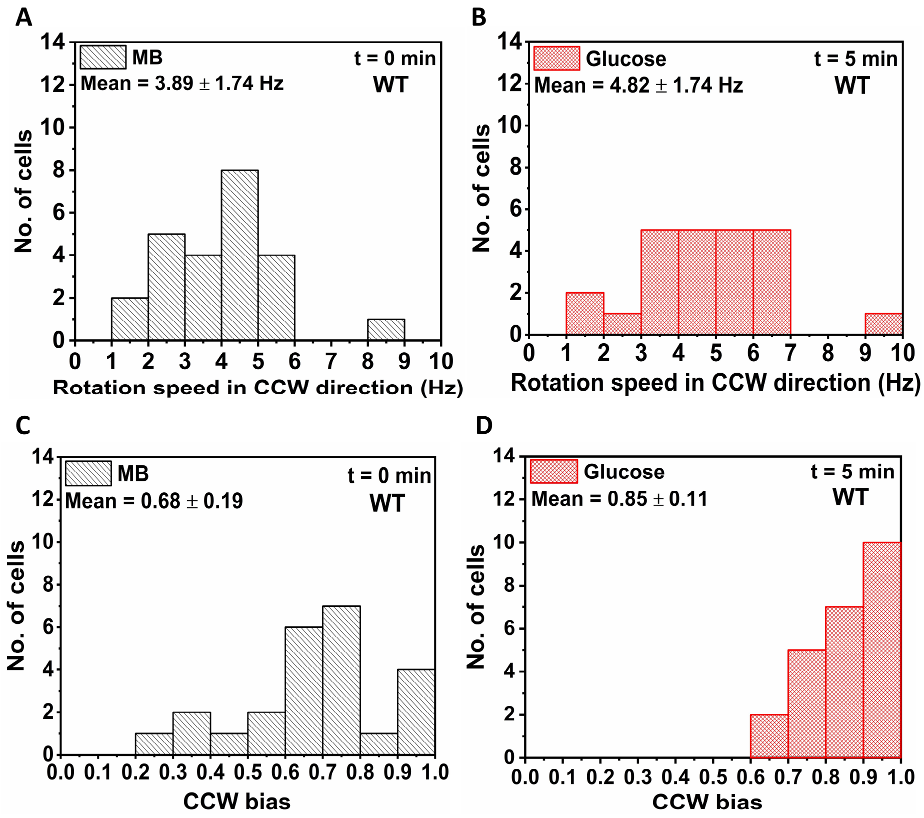


FIG. S5. Response of WT after 5 min of exposure to 1000 $\mu$M of glucose. Rotation frequency of 24 cells in (A) MB, and (B) when exposed to 1000 $\mu$M glucose. The corresponding CCW directional bias are given in (C) and (D). Weighted average means are calculated for cell population obtained from four independent experiments. All values are significantly different from MB at $p < 0.001$ computed by paired t-test.

Figure S6 presents the normalized rotation speed and the CCW bias for 5 cells after exposure to 1000 $\mu$M of glucose and 2Dg. As expected, the rotation speed and the CCW bias increase with time. While the CCW bias in case of 2Dg returned to its pre-stimulus value, the same was not observed for glucose even after 15 min exposure. The variation of the measured CCW values among cells are presented in Figure S6B, D.

Figure S7 shows the distribution of rotational frequency in CCW direction for at least 20 cells each of WT, $\Delta trg$ and $\Delta ptsI$ cells measured in MB, and 1000 $\mu$M of glucose and

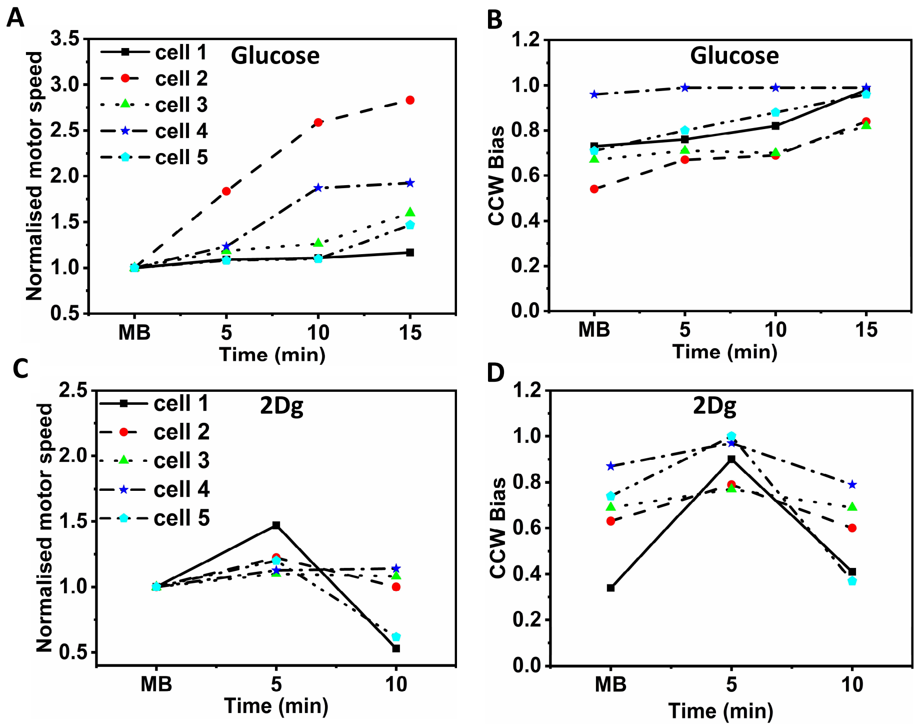


FIG. S6. Rotation speed normalized with respect to the pre-stimulus value and the CCW bias of WT at discrete time points in the presence of (A,B) 1000 $\mu$M glucose and, (C,D) 1000 $\mu$M 2Dg.

2Dg. The mean rotational speed of WT in MB was measured to be about 4.14±1.65 Hz (Fig S7A), which compares well with the previous studies [7] . Experiments with glucose resulted in a rotation speed of 5.95±1.69 Hz at 15 min (Fig S7B), an increase of 44 % in motor speed compared to that observed in MB. At the intermediate time of 5 min, motor speed of 4.82±1.74 Hz (Fig S5B) was obtained which corresponds to about 25 % increase as compared to MB, which is inline with the trend observed for run speeds. In comparison, an increase of 27 % (5.28±1.74 Hz) was observed at 5 min in the presence of 2Dg (Fig S7C). At longer times, the rotation speed decreased significantly, as also observed in case of run speed, due to the lack of energy source. These results clearly demonstrate that sensing alone can enhance the motor speed leading to an increase in run speed. These measurements also yield CCW bias for the three cases. A CCW bias of 0.90±0.09 was observed for glucose at 15 min (Fig S8B) and 0.78±0.16 for 2Dg at 5 min (Fig S8C), compared to 0.68±0.16 in MB (Fig S8A).

Tethered cell experiments were also performed with $\Delta trg$ mutant strain.Figure S7D-

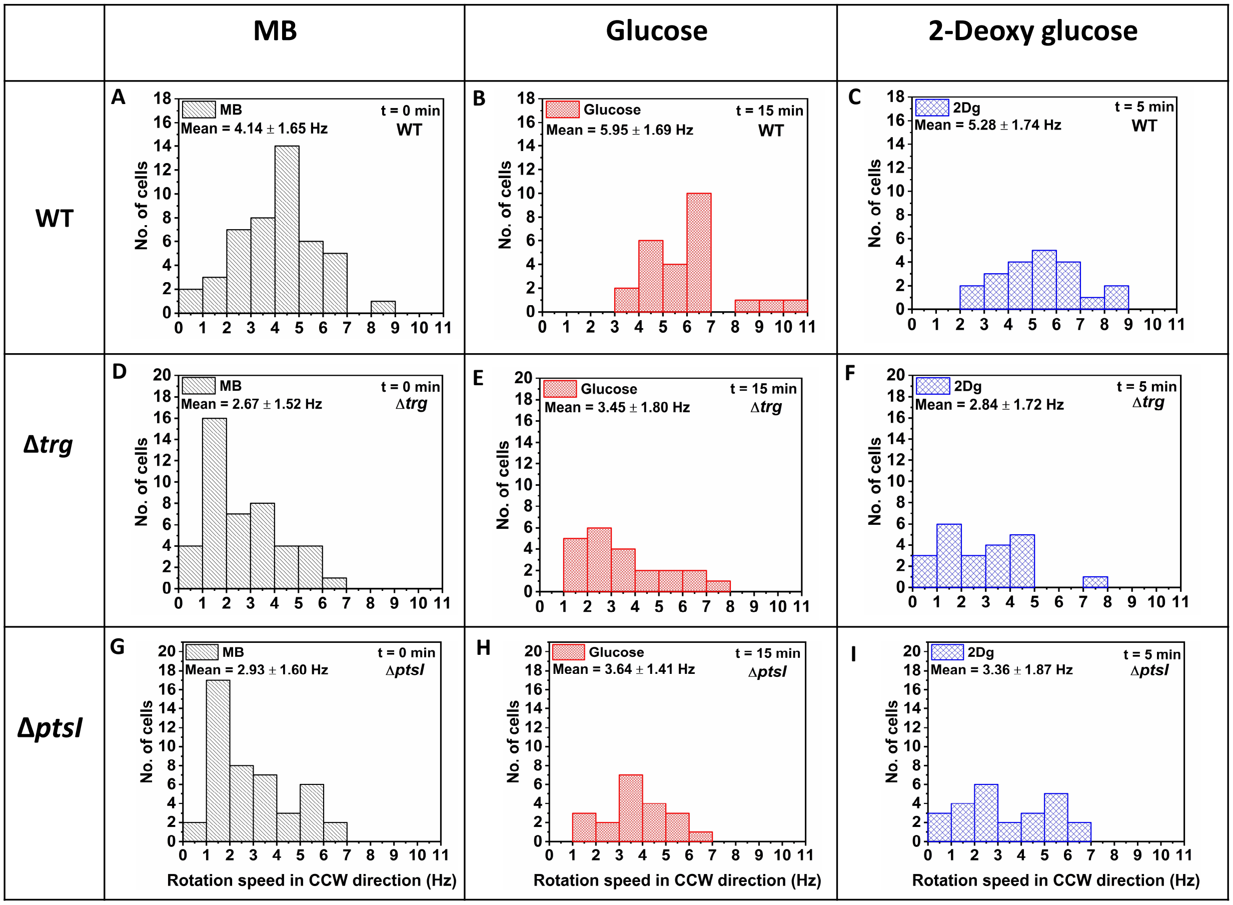


FIG. S7. Tethered cell experiment with the (A-C) WT, (D-F) $\Delta trg$ and (G-I) $\Delta ptsI$. Measured rotation speed of individual cells were first obtained in MB and then measured again for the same cells exposed to 1000 $\mu$M of glucose and 2Dg. At least 20 such paired cells from four independent experiments were analyzed to determine the population behavior. The data for MB is obtained by combining the measurements from the pre-stimulus state of both glucose and 2Dg experiments. Values for glucose and 2Dg are obtained after 15 min and 5 min of exposure, respectively. All values are significantly different from MB at $p < 0.05$ computed by paired t-test except the bias and motor speed change from MB to 2Dg for $\Delta ptsI$ strain, which was statistically insignificant.

F represents the distribution of rotation speed in CCW direction in MB, glucose and 2Dg respectively. It can be clearly seen that the mean rotation speed increased by 30% (3.45$\pm$1.80 Hz) in glucose as compared to MB, whereas no significant variation was observed when the cells were exposed to 2Dg. This reiterates the role of Trg receptor in modulating the motor speed. The observed increase in rotation speed in the presence of glucose is caused by the glucose PTS uptake mechanism. The CCW bias mimicked the trend observed for the rotation speed, wherein CCW bias increased by 13% in glucose related to MB, with no change in 2Dg (Fig S8D-F).

A similar exercise performed with $\Delta ptsI$ mutant strain (Fig S7G-I) showed a 24% increase in rotation speed in the presence of glucose compared to MB. However, the small increase (16%) observed in 2Dg was not statistically significant ($p > 0.05$). Further, the change observed in the $\Delta ptsI$ strain compared to $\Delta trg$ strain was lower. A similar trend was observed for the CCW bias (Fig S8G-I).

Figure S8 presents the CCW bias for the WT, $\Delta trg$ and $\Delta ptsI$ cells measured in MB, and 1000 $\mu$M of glucose and 2Dg. In case of glucose, the CCW bias increases with time and remains higher than the pre-stimulus value for all strains even after 15 min. In case of 2Dg, while the bias returns to the pre-stimulus value within 5 min for both $\Delta trg$ and $\Delta pstI$, it takes longer (between 5-10 min) in case of WT (see Fig S6).

Figure S9 presents the variance of the rotation rate for increasing number of revolutions recorded for a single WT cell exposed to 1000 $\mu$M of glucose, and 2Dg. At a fixed time, the variance increases with number of revolutions but the magnitude of the slope decreases with time in the presence of glucose. However, in case of 2Dg, the slope decreases initially and then increases after 5 min. Decreasing variance indicates increasing number of torque-generating units leading to a smoother rotation of the motor. These results are consistent with the behavior observed both for single motor and a population.

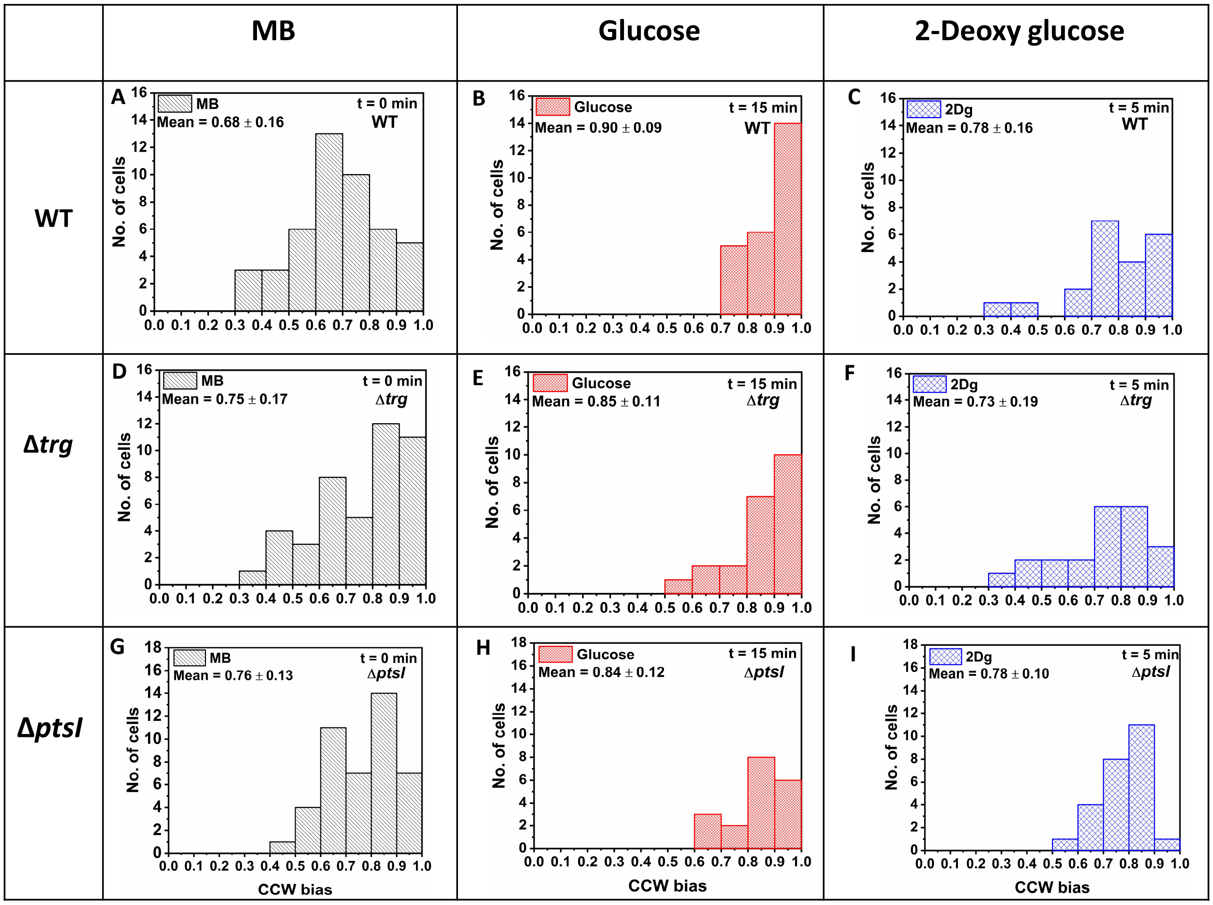


FIG. S8. CCW bias for the WT, $\Delta trg$ and $\Delta ptsI$ cells measured in MB, and 1000 $\mu$M of glucose and 2Dg. At least 20 such paired cells from four independent experiments were analyzed to determine the population behavior. The data for MB is obtained by combining the measurements from the pre-stimulus state of both glucose and 2Dg experiments. Values for glucose and 2Dg are obtained after 15 min and 5 min of exposure, respectively. All values are significantly different from MB at $p < 0.05$ computed by paired t-test except the change in bias from MB to 2Dg, which was statically insignificant.

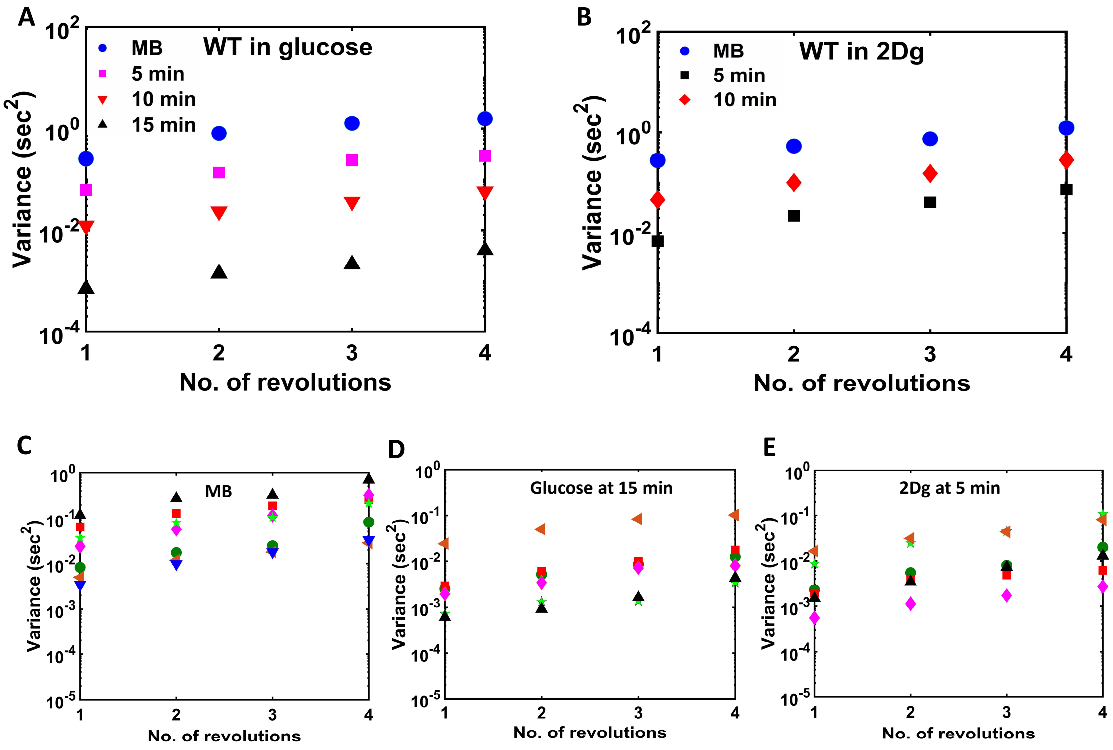


FIG. S9. Variance of rotation rate for different number of revolutions determined at different times after exposing a single WT cell to 1000 $\mu$M of (A) glucose and, (B) 2Dg. As compared to (C) MB (7 cells), the variance decreased on introduction of (D) glucose (after 15 min) (6 cells) and (E) 2Dg (5 min) (6 cells).
